# Rationally inattentive intertemporal choice

**DOI:** 10.1101/680652

**Authors:** Samuel J. Gershman, Rahul Bhui

## Abstract

Discounting of future rewards is traditionally interpreted as evidence for an intrinsic preference in favor of sooner rewards. However, temporal discounting can also arise from internal uncertainty in value representations of future events, if one assumes that noisy mental simulations of the future are rationally combined with prior beliefs. Here, we further develop this idea by considering how simulation noise may be adaptively modulated by task demands, based on principles of rational inattention. We show how the optimal allocation of mental effort can give rise to the magnitude effect in intertemporal choice. In a re-analysis of two prior data sets, and in a new experiment, we reveal several behavioral signatures of this novel theoretical account, tying choice stochasticity to the magnitude effect. We conclude that some aspects of temporal discounting may result from a cognitively plausible adaptive response to the costs of information processing.

## Introduction

The preference for sooner over later rewards is traditionally interpreted as an intrinsic decline in value as outcomes recede into the future. However, recent evidence suggests an alternative (though not mutually exclusive) viewpoint: temporal discounting could arise from *internal uncertainty* in value representations of future rewards. Imagining the future allows an agent to immediately experience anticipated outcomes, helping them to delay gratification, but this prospection may lose its impact when mental simulations are noisy. A number of influential studies show that patience is enhanced by treatments that may be thought of as increasing the precision of mental simulation. For example, discounting is attenuated when people are asked to imagine spending future rewards [1], when they imagine future outcomes in greater detail [2], and when episodic tags are provided to facilitate such imagination [3].

Thus, even if agents with imperfect foresight intrinsically valued delayed rewards just as much as immediate rewards, the myopic nature of their prospection could reduce the impact of the future. This idea has recently been formalized in a Bayesian model of discounting [4], in which an agent observes noisy simulations of future value and applies Bayes’ rule to obtain a posterior estimate. Assuming simulations become noisier the further they reach into the future, the agent increasingly relies on their prior beliefs and discounts the reward value. Gabaix and Laibson [4] showed how this can lead to hyperbolic discounting while accommodating the effects of experience on intertemporal choice tasks. However, this analysis is predicated on a fixed relationship between reward delay and simulation noise, though there is reason to think that the relationship is not fixed. We extend this perspective by considering how the degree of simulation noise may be adaptively controlled, and propose how such a mechanism contributes to the well-known *magnitude effect* in intertemporal choice—the finding that people are disproportionately more patient when judging high-value outcomes [5, 6, 7, 8, 9].

According to our theory, vivid prospection can help agents to delay gratification, but this comes at a cost. Making simulations more precise requires mental effort, and this effort may only be invoked if its benefits outweigh its costs. We formalize this intuition in terms of *rate-distortion theory*, an information-theoretic framework for modeling the optimal level of internal uncertainty ([10]; see also [11, 12, 13]). Richer simulations are cognitively costly, and therefore a decision maker must make a trade-off involving precision and effort. Most relevant in the present context, larger rewards may be more important to evaluate carefully. In this case, greater magnitudes would be simulated more precisely and, in light of the above argument, would engender more patience. The model thus implies a direct connection between stochasticity and discounting.

Our theory is consistent with several lines of psychological evidence. Mental representations of events farther in the future generally contain fewer sensory and contextual details than those closer in time [14, 15]. Future events are imagined with greater vividness when cued by more rewarding stimuli [16], and people produce longer lists of thoughts when prompted to evaluate higher magnitude intertemporal choices [17]. Moreover, when people are asked to write down justifications for their choices, patience is enhanced specifically for lower magnitude rewards, as if cognitive control is already being exerted at higher magnitudes [18].

In what follows, we investigate the behavioral implications of this theory. We show how it qualitatively accounts for several empirical findings pertaining to the magnitude effect, quantitatively improves model fit in a large existing data set, and accurately predicts patterns of discounting and stochasticity in a new experiment. These results help sharpen our understanding of the relationship between patience, reward, and mental effort.

## Results

### A Bayesian model of as-if temporal discounting

In this section, we first describe the Bayesian model of discounting developed by Gabaix and Laibson [4]. In the next section, we extend this analysis by endogenizing the simulation noise variance using a rational inattention analysis.

Following Gabaix and Laibson [4], we model an agent who is faced with a choice between several rewards that occur at some time in the future. For ease of exposition, we will consider a single reward r_t_ delivered after delay *t*, whose “true value” is denoted by u. We assume that this value is drawn from a Gaussian distribution with mean *μ* and variance 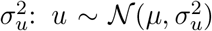. We further assume that the agent does not directly observe u, but instead observes a noisy signal 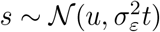 generated by some form of mental simulation.

Noise arises from the agent’s limited ability to simulate the event’s future value. Gabaix and Laibson [4] assumed that the variance increases linearly with the delay because events farther in the future are harder to simulate. Combined with the assumption that the prior mean μ is 0 (which we suppose for the remainder of the paper), this leads to the following expression for the posterior mean:

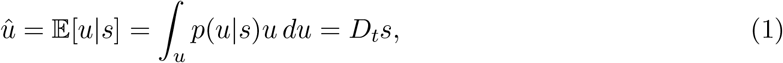

where *p*(*s|u*) is the posterior, computed using Bayes’ rule:

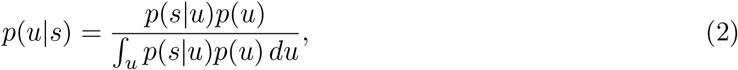

with likelihood *p*(*s|u*) and prior *p*(*u*) as defined above. The term *D_t_* expresses an “as-if” hyperbolic discount function:

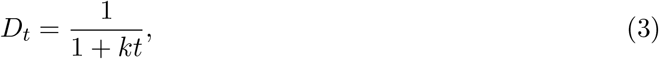

with the as-if discount rate *k* given by:

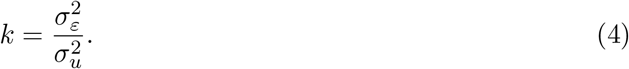

The discount function is “as-if” because the agent in fact has a neutral time preference, but chooses in accordance with hyperbolic discounting, one of the most broadly supported models of intertemporal choice (see [19] for a review). Figure 1 illustrates how Bayesian inference in this model produces temporal discounting.

**Figure 1:**
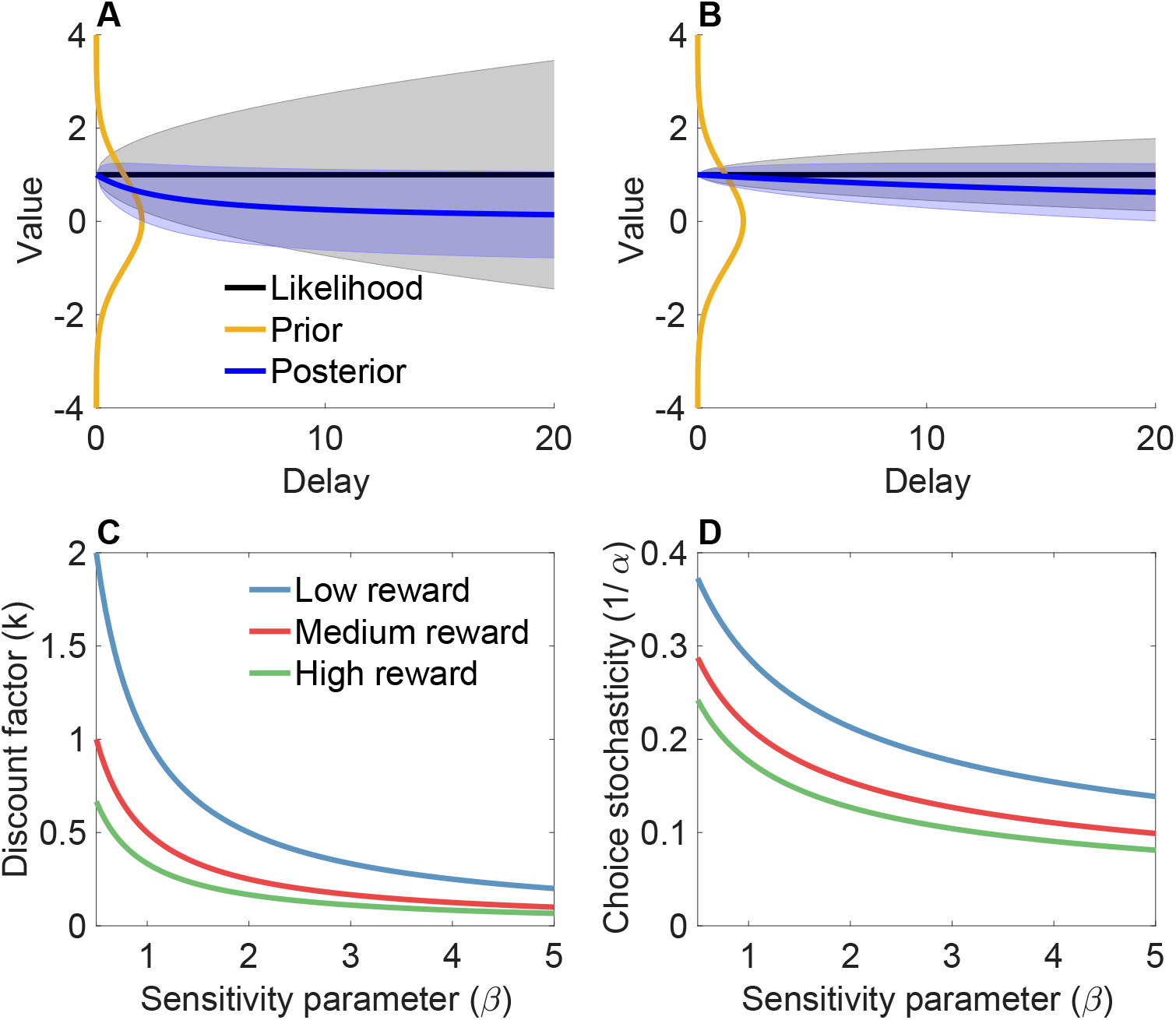
(A) Illustration of discounting as Bayesian inference. Mental simulation generates a noisy value signal (*s_t_* = 1), which is combined with a prior distribution to form a posterior distribution over value. The black line denotes the true underlying value *u*, and the shaded region shows the standard deviation of the noise corrupting the value at each possible delay *t* (in arbitrary units). Similarly, the blue line is the posterior mean, and the shaded region is the posterior standard deviation. Because the standard deviation of *s_t_* (simulation noise) increases with delay, the posterior mean is pulled more strongly towards the prior mean for longer delays. (B) Same simulation as in (A) but with a smaller signal variance, demonstrating a reduction in discounting. (C) Discount factor under the rational inattention model as a function of the sensitivity parameter (*β*) and reward magnitude. (D) Choice stochasticity under the rational inattention model as a function of the sensitivity parameter (*β*) and reward magnitude.

The estimated value of a reward will be regularized towards the mean *μ* (0 in this case). The strength of this regularization depends on *k*, which can be thought of as an inverse signal-to-noise ratio. Intuitively, when the simulation noise variance 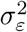 is large relative to the prior variance 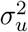, the simulations are less reliable and the agent will rely more on their prior, whereas when the simulation noise variance is relatively small then agent will rely more on their simulations.

Because we (the experimenters) cannot directly observe the signal *s*, we use the objective reward *r_t_* as a proxy. This allows us to link the model directly to experimentally observable variables. We note, however, that this assumption may generate erroneous inferences. For example, we may misinterpret the effects of model misspecification in terms of simulation noise.

### Rational inattention

The Gabaix and Laibson [4] analysis assumed that the agent has a fixed simulation noise variance. Here we develop the idea that the simulation noise variance is determined by the agent’s “attention” to the signal. Intuitively, an agent can improve the reliability of their mental simulations by exerting cognitive effort (i.e., attending more), but pays a cost for this effort.

We approach this problem through the lens of rate-distortion theory [10]. Rate-distortion theory offers us a principled way to study the optimal precision of internal representations, formalized using information theory. As such it has been fruitfully applied to human cognition in domains such as perceptual judgment and working memory [20], and its close relative, rational inattention [11, 21], has been used to analyze a variety of economic problems [12, 13]. In this framework, the agent is modeled as a communication channel that takes as input the signal and outputs an estimate of the value. The agent can select the design of the channel subject to a constraint on the information rate of the channel (the number of bits that can be communicated per signal).

In this case, we define a family of channels parametrized by the simulation noise scaling parameter, 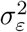. The optimization problem is to select the value of 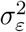 that minimizes the expectation of a squared error distortion (aka loss) function that quantifies the cost of estimation error. As shown in the Methods section, the optimal simulation noise parameter under some assumptions is given by:

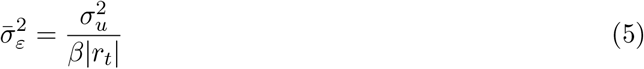

where *β* > 0 is a “sensitivity” parameter that governs the link between information rate and magnitude. As *β* increases, the rate becomes increasingly sensitive to variations in reward and delay. Plugging this into `Eq. 4 yields the optimal discount parameter:

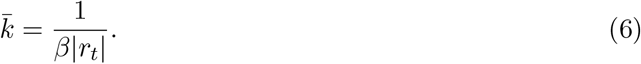

Thus, the rate-distortion framework can lead us to a model that captures the magnitude effect (inverse relation between discount factor and reward magnitude; Figure 1C). As shown in the Methods, the model also predicts a choice stochasticity magnitude effect: choices should become less stochastic as magnitude increases (Figure 1D). This arises in the model because choice stochasticity is partially driven by simulation noise, which should decrease with reward magnitude.

### Applications to prior experimental results

In this section, we explore the empirical implications of the rational inattention analysis. We begin by examining experimental data collected by Ballard and colleagues [18], in which subjects reported their indifference point between an immediate and delayed reward. The reward magnitude was manipulated across subjects (see Methods for more details). In addition, some subjects were assigned to a “justification” condition in which they were asked to explicitly justify their choices. Ballard and colleagues hypothesized that the magnitude effect arises from increased self-control in response to large magnitudes, and reasoned that justification would elevate the ceiling on selfcontrol. In the language of rational inattention, we interpret justification as prompting increased allocation of cognitive resources to prospective simulations. This hypothesis can be formalized by increasing the *β* parameter in the justification condition compared to the no justification condition.

Five predictions follow from this hypothesis, all of which are confirmed in Figure 2, and quantified by a regression with regressors for justification (no justification coded as +1, justification coded as 0), magnitude, and the interaction between justification and magnitude (negative coefficient indicates a *reduced* justification effect for larger magnitudes). For all of the following analyses, we report bootstrapped 95% confidence intervals. First, the average discount factor *k* should be larger in the no justification condition, because *k* decreases monotonically with *β* (regression coefficient for the main effect of justification: CI = [0.064, 0.155]). Second, the justification effect should diminish with magnitude, because *dk/dβ* is a concave function of |*r*| (regression coefficient for the interaction: CI = [−0.036, −0.015]).

**Figure 2:**
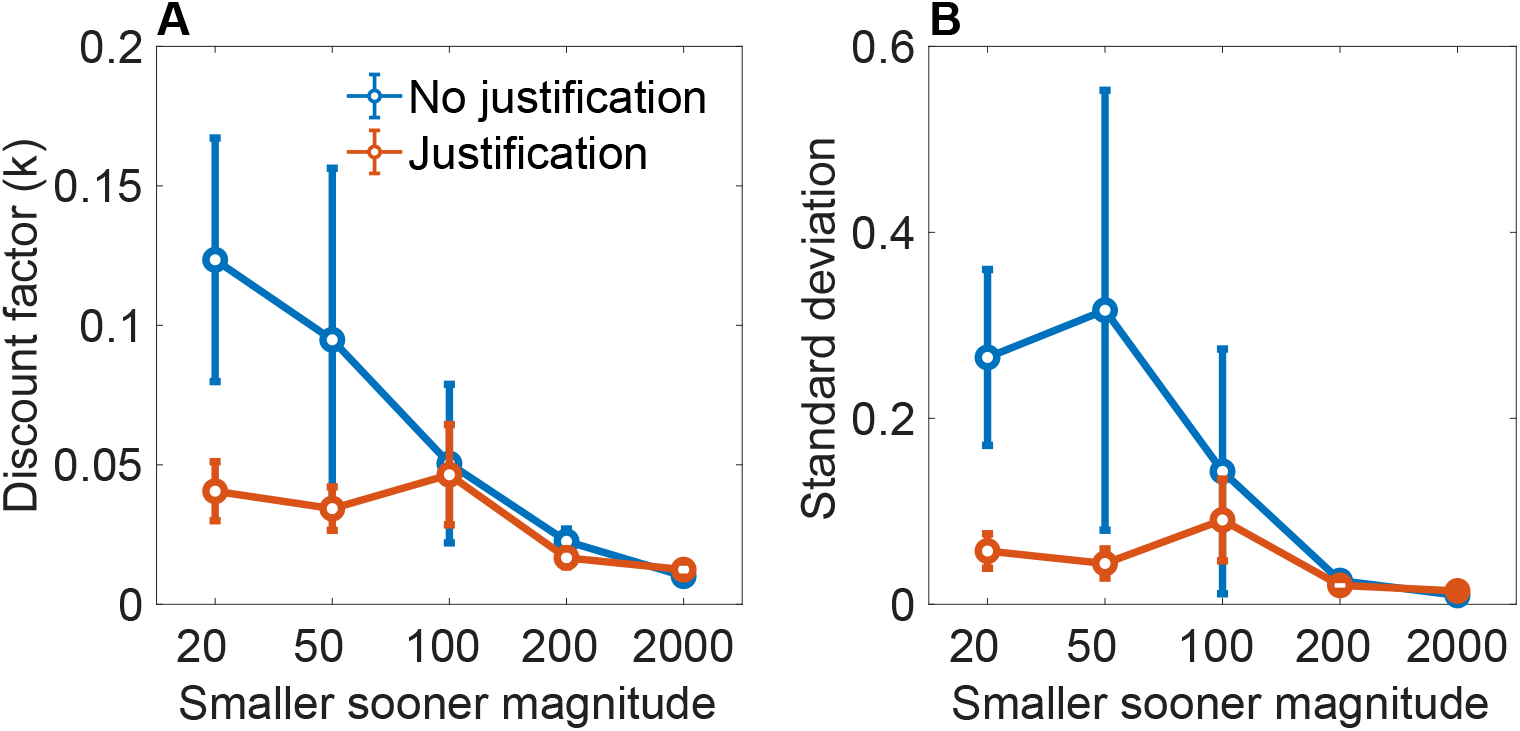
(A) Mean discount factor as a function of magnitude and justification condition. (B) Standard deviation of the discount factor as a function of magnitude and justification condition. Data from [18]. Error bars show 95% confidence intervals.

The next three predictions are distinctive of our theory, and pertain to the variability of *k*, which we quantify by the standard deviation. The third prediction is that the standard deviation of *k* should be higher for small magnitudes (i.e., a magnitude effect for response variability; regression coefficient for the main effect of magnitude: CI = [−0.016, −0.008]). The fourth prediction is that the standard deviation should be lower in the justification condition, because response variability decreases with *β* (regression coefficient for the main effect of justification: CI = [0.203, 0.501]). The fifth prediction is that the justification effect for response variability should diminish with magnitude, provided that *β*|*r*| is sufficiently large relative to the delay (regression coefficient for the interaction effect: CI = [−0.109, −0.045]).

The Ballard data set confirms several predictions qualitatively, but is ill-suited to confirming quantitative predictions because each subject only saw a single experimental condition. To quantitatively assess the validity of our model, we re-analyzed a large data set (*N* = 1284) of intertemporal choices collected by Chavez and colleagues [22]. Each subject in this study was presented with the same set of 27 choices, taken from [7]. The rewards for both options and the delay for the larger-later option varied across trials, while the delay for the smaller-sooner option was held fixed at 0 days.

We compared our rational inattention model with several alternatives using random-effects Bayesian model selection (see Methods). In particular, we compared the full rational model (R2) to a variant (R1) which uses the optimal discount factor but treats the inverse temperature *α* as a free parameter. We also compared against standard quasi-hyperbolic discounting (QH; [23]), and several variations of hyperbolic discounting, including the basic functional form (H0), and generalized versions that incorporate magnitude-dependent discounting and choice stochasticity (H1-H3; [24]). We used the protected exceedance probability (PXP) as a measure of model evidence. The PXP measures the probability that a particular model is more frequent in the population than all the other models under consideration, adjusting for the probability of differences arising from chance.

We found that the full rational inattention model (R2) was decisively favored (PXP > 0.99). Among the 4 variants of hyperbolic discounting, H3 was favored. We used this model to assess the qualitative predictions of the rational inattention theory (note that the rational inattention theory *assumes* discounting and choice stochasticity magnitude effects, so it cannot be used to falsify these predictions). Consistent with the theory’s predictions, the magnitude scaling parameter for inverse temperature (*m_α_*) was significantly greater than 0 [t(1283) = 7.47, *p* < 0.0001], indicating that choice stochasticity decreases with reward magnitude, whereas the magnitude scaling parameter for discounting (*m_k_*) was significantly less than 0 [t(1283) = 15.42,p < 0.0001], indicating that myopia decreases with reward magnitude (Figure 3A). Finally, we observed that the two magnitude scaling effects are negatively correlated (*r* = −0.21,p < 0.0001; Figure 3B), consistent with the rational inattention model’s predictions. Thus, the data support the theory both qualitatively and quantitatively.

**Figure 3:**
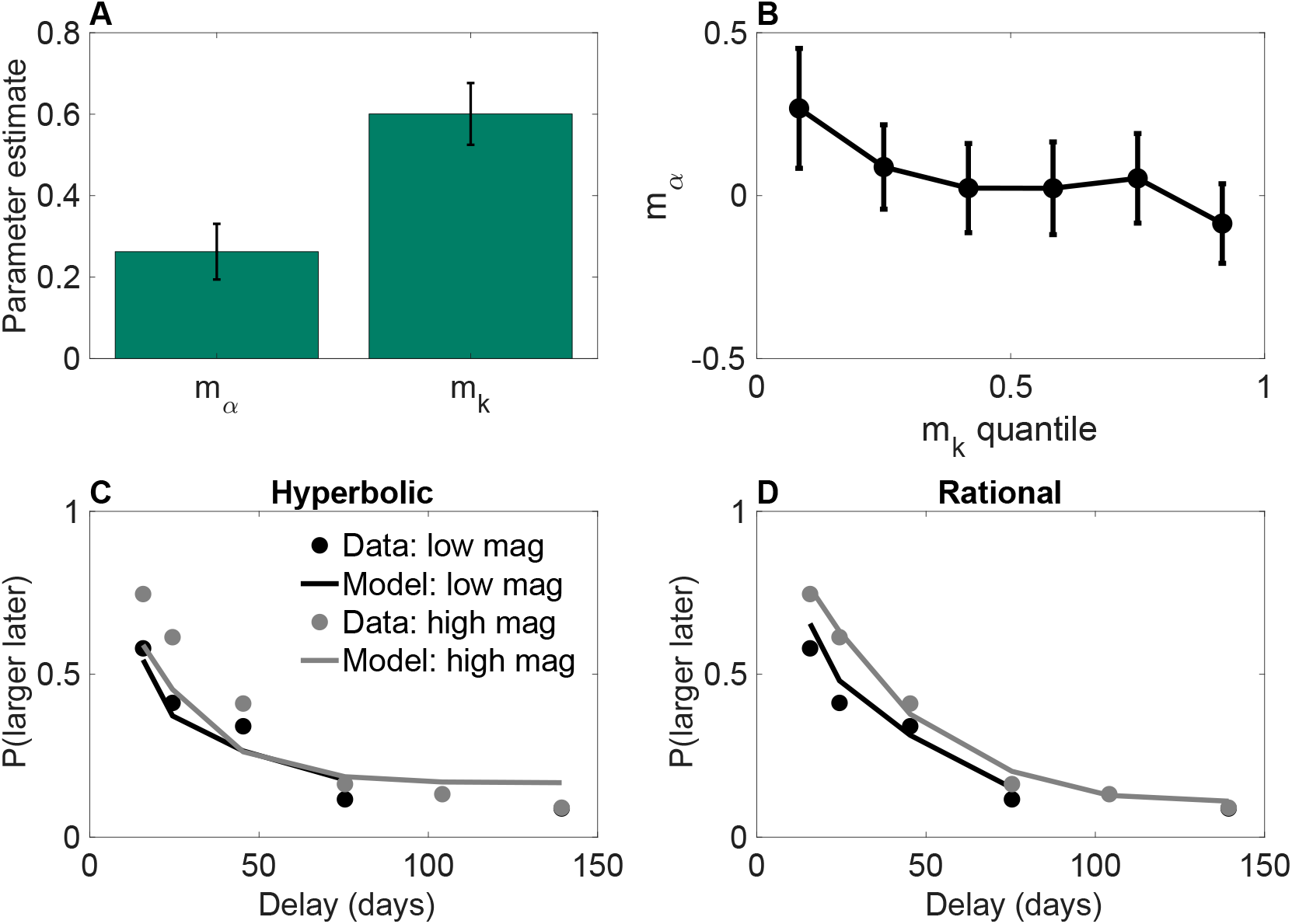
(A) Magnitude scaling parameter estimates for the 3-parameter hyperbolic discounting model (H3). Error bars show 95% confidence intervals. (B) The magnitude scaling parameter for choice stochasticity declines as a function of the magnitude scaling parameter for discounting. (C) Choice probabilities as a function of delay for low and high variance conditions, with theoretical functions obtained from the standard hyperbolic discounting model (H0). (D) Same as (C), but with theoretical functions obtained from the rational discounting model (R2).

To further support the rational inattention model, we compared the psychometric functions of standard hyperbolic discounting (H0) and the full rational inattention model (R2), finding that choice probabilities were much better fit by R2, despite having fewer parameters (Figure 3C-D).

### The effect of reward variance on discounting and choice stochasticity

The rational inattention model predicts that the choice stochasticity magnitude effect should decrease with reward variance, because the noisy simulations become increasingly down-weighted as the reward variance increases, and this down-weighting interacts multiplicatively with the reward magnitude. The model also predicts that there should be no effect of reward variance on the discounting magnitude effect. We tested these predictions in a new experiment (*N* = 221) in which the reward variance was manipulated while holding the mean and range of rewards fixed.

To evaluate the variance predictions, we fit the same models described above to the choice data. In this case, the model with the strongest support (PXP = 0.61) was H3 (hyperbolic discounting with magnitude-dependent discounting and choice stochasticity). The key parameter estimates are shown in Figure 4, broken down by variance condition. Replicating our prior results with the Chavez data set, we found a significant discounting magnitude effect [*m_k_* < 0: *t*(220) = 7.16,*p* < 0.0001] and a significant choice stochasticity magnitude effect [*m_α_* > 0: *t*(219) = 9.95,*p* < 0.0001] when collapsing across conditions. Critically, the choice stochasticity magnitude effect was significantly lower in the high variance condition, [*t*(219) = 2.22,*p* < 0.05], whereas there was no effect of variance on the discounting magnitude effect (p = 0.84). Using a Bayesian t-test with a scaled JZS prior [25], we found a posterior probability > 0.99 favoring the null hypothesis that variance does not modulate the discounting magnitude effect. These results collectively provide evidence consistent with our rational inattention model.

**Figure 4:**
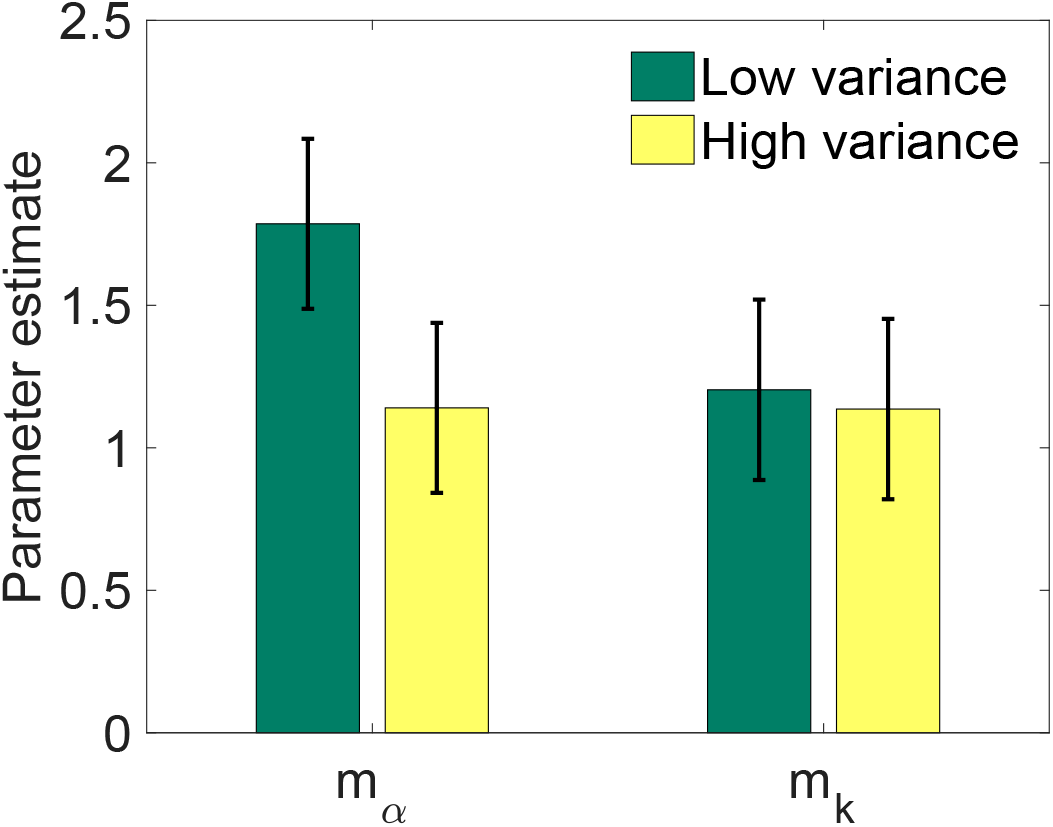
Magnitude scaling parameter estimates for the 3-parameter hyperbolic discounting model (H3), aggregated separately for low and high variance conditions. Error bars show 95% confidence intervals.

## Discussion

Temporal discounting may stem partly from the inability of decision makers to perfectly simulate future outcomes [26]. In this paper, we develop a theoretical account of prominent regularities in intertemporal choice, based on the idea that mental simulation of the future is noisy but controllable. Our approach connects the Bayesian model of discounting from [4] with the information-theoretic framework of rate-distortion theory [10] (see [20] for an overview of rate-distortion theory applications to human perception; see [11, 12, 13] for closely related economic applications of rational inattention). Supposing the prospective value of a reward becomes noisy when it is internally projected into the future, Bayesian agents should compensate for this uncertainty by relying more heavily on their prior beliefs—if priors are centered near zero, this leads to discounting of value. However, the degree of noise in the simulation may be controlled by the agent, at a cost. If it is more important to accurately evaluate larger rewards, the agent should spend extra mental effort to make their simulations more precise when dealing with greater magnitudes. This mechanism could lead to reduced temporal discounting when dealing with large rewards, a commonly observed phenomenon known as the magnitude effect. Our model can also account for how reward magnitude and contextual variability are simultaneously related to stochasticity in choice, which we validate in the re-analysis of two data sets and a new experiment.

Note that the uncertainty we are dealing with is *internal*. This contrasts with theories of discounting based on objective risk in the arrival of rewards (e.g., [27, 28]). In the present framework, discounting can occur even when the decision maker has no innate preference for earlier rewards and there is no extrinsic risk. Of course, all of these pathways are not mutually exclusive, and we do not claim the others are inconsequential. Our goal is rather to clearly describe how apparent anomalies of intertemporal choice could arise from a cognitively plausible adaptive response to limits on information processing.

Our proposal is supported by a range of neural and behavioral evidence. Psychologically speaking, the allocation of attention in our framework (and what Gabaix and Laibson refer to as “mental effort”) may manifest as *cognitive control*—the set of mechanisms required to pursue a goal in the face of distractions and competing responses. It has been argued that the exertion of cognitive control depends on its expected value, the combination of its effort costs and payoff benefits in a given task, and that this plays a role in many decisions including intertemporal choices [29]. Future events have been found to be imagined with greater vividness when cued by more rewarding stimuli [16], and people list a greater number of thoughts when prompted to evaluate higher magnitude intertemporal choices [17]. Moreover, when people are asked to explicitly justify their choices, they exhibit more patience specifically for lower magnitude rewards, as if they have already hit a ceiling for higher magnitudes [18]. Our model formally draws out the implications of this cost-benefit logic, providing a high-level normative perspective that complements more mechanistic analyses of cognition and discounting (e.g., [30, 31]).

From a neuroscientific perspective, the exertion of cognitive control is known to rely on a network of regions in prefrontal cortex, which some studies have linked directly to temporal discounting [32, 1]. Shenhav and colleagues [29] have proposed that the expected value of control is computed by connected regions and guides the investment of attention into each task, while Ballard and colleagues [18] demonstrated that frontal executive-control areas of the brain are particularly engaged in challenging intertemporal choices with high-magnitude rewards. Moreover, disruption of activity in such areas via transcranial magnetic stimulation reduces the magnitude effect [33]. Taken together, these studies indicate that the brain adaptively modulates simulation noise and this plays a meaningful role in temporal discounting.

Another perspective from neuroscience is provided by studies of patients with Parkinson’s disease, who are known to have systemically low levels of dopamine. Foerde and colleagues [34] observed that patients on medication (with putatively higher dopamine levels) exhibit both more patience (higher estimated values of the *k* parameter) and a weaker magnitude effect compared to patients off medication. Both of these findings are consistent with the idea that higher levels of dopamine correspond to higher values of the sensitivity parameter *β*. Higher sensitivity means that reward will induce a greater willingness to exert cognitive effort, which in this case means reducing simulation noise and thereby reducing discounting. At the same time, increases in sensitivity will actually make the magnitude effect smaller, because of the concave relationship between the discount parameter and reward magnitude. Our interpretation of dopamine in terms of sensitivity is consistent with other work on Parkinson’s patients showing that high levels of dopamine produce greater reward sensitivity [35, 36]. More broadly, it has been suggested that dopamine may control allocation of cognitive effort [37]. We conjecture that dopamine may play a specific role in mediating the relationship between reward and information rate, but further research will be required to directly test this hypothesis.

An important limitation of our experimental study was the hypothetical nature of choices made by subjects, a design element prompted by the impracticality of payment at the lengthy delays needed to precisely estimate discount rates. Many of the classic (e.g., [5, 6, 38]) and modern (e.g., [18, 39]) studies of the magnitude effect are not incentive compatible, for the same reason. A recent survey has argued that comparison between incentive compatible and incompatible designs typically yield the same results for studies of intertemporal choice [40]. For example, Bickel and colleagues have found that discount rates are highly correlated across real and hypothetical rewards [9], as are their neural responses [41]. Moreover, according to our analysis (following Gabaix and Laibson), all decisions involve some future simulation, with the difference resting in the degree of simulation noise. Thus, although incentive compatibility is an important criterion towards which to strive in decision making studies, practical and theoretical considerations render it less applicable to the experimental questions pursued here.

Finally, while our theory naturally captures a number of empirical phenomena surrounding the magnitude effect, future work may examine what other observations might be accommodated under other assumptions. For instance, people seem to savor and dread future outcomes [42], which could lead people to prefer early resolution of losses, and the magnitude effect has been found to reverse in the loss domain [17]. The Bayesian discounting model cannot account for this inverted pattern, as it implies that deferred losses should be treated better than immediate ones. Nonetheless, there might be adaptive value in anticipation if it could facilitate planning and decision making [43]. A more formal account of the costs and benefits involved may help predict when people will channel energy into such anticipatory thoughts.

## Methods

### Derivation of optimal precision

In order to derive the optimal precision 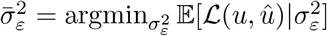, the expected quadratic loss is computed as follows. Conditioning on *u* and *s*,

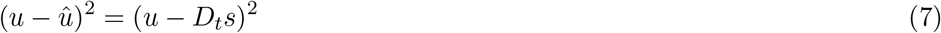

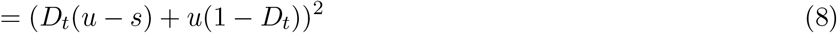

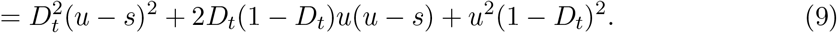

Taking the expectation over *p*(*s|u*), and subsequently over *p*(*u*),

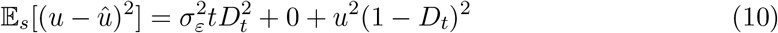

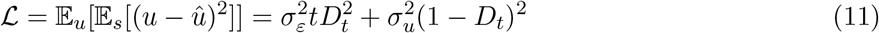

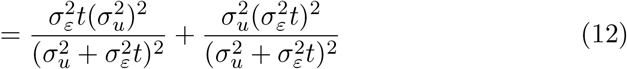

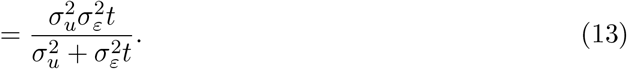

We then plug this into the rate-distortion function for a Gaussian source (which reflects the rate-distortion frontier, that is, the minimal achievable information rate for a given distortion level, or equivalently the minimal achievable distortion for a given rate):

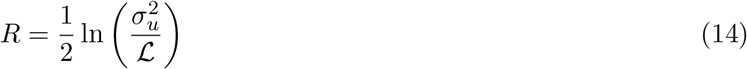

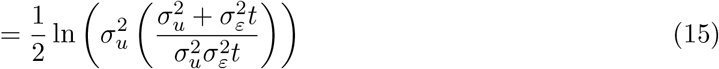

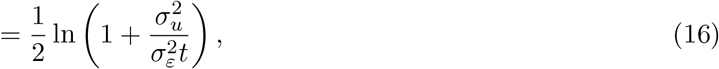

which can be rearranged to yield the optimal precision:

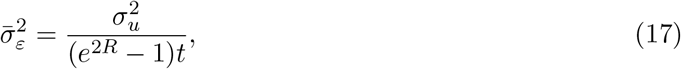

where *R* is the information rate constraint in nats (i.e., units of information in base *e*).

We impose an additional constraint on this formulation, by assuming that information rate increases with reward magnitude (greater incentive to expend cognitive resources) and decreases with delay (simulation of distal events is more cognitively demanding):

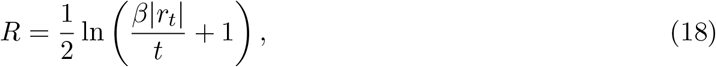

where *β* > 0 is a “sensitivity” parameter that governs the relationship between rate, magnitude and delay. As *β* increases, the rate becomes increasingly sensitive to variations in reward and delay. The constraint follows in the spirit of Gabaix and Laibson’s framework, reflecting a costand-benefit perspective on their baseline assumptions. The greater cost of simulating more distal events parallels their supposition of greater noise for projections extending farther into the future. Plugging R into `Eq. 17 yields the optimal simulation noise parameter:

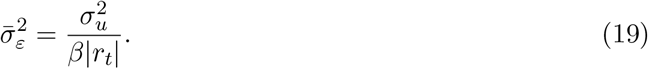

Note that although the optimal simulation noise variance in `Eq. 17 depends on t, this dependence disappears when we use the rate constraint specified in `Eq. 18.

We can draw out further implications of this model by connecting it to choice behavior. Let us assume, in the simplest case, that the agent deterministically chooses the option with highest estimated value. In this case, all stochasticity in choice behavior is driven by stochasticity in the agent’s simulation process. Marginalizing over these noisy simulations, the choice probability for a standard two-alternative choice (early vs. late) is given by:

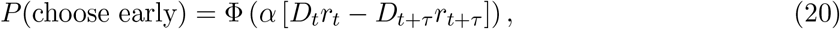

where *ϕ* is the standard Gaussian cumulative density function, *τ* is the difference in delay between early (*r_t_*) and late (*r_t+τ_*) options, and

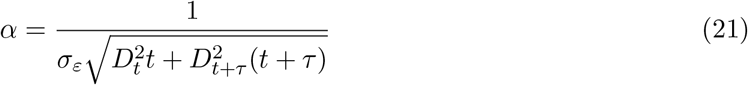

is an “inverse temperature” parameter controlling the degree of choice stochasticity (smaller values of *α* produce greater stochasticity). In the case where the early option is immediate (i.e., *t* = 0), as in many studies of discounting, this simplifies to:

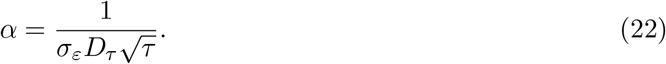

Plugging in the optimal simulation noise parameter gives:

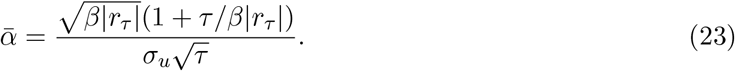

One can show that

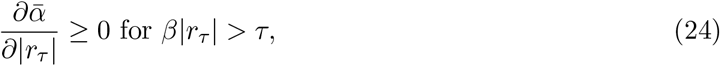

which means that for sufficiently large rewards and sufficiently short delays, the model predicts a choice stochasticity magnitude effect: as reward magnitude gets larger, choice stochasticity should get smaller. One can also show that

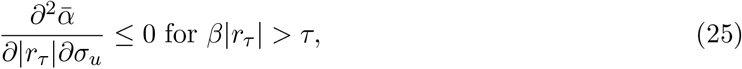

which means that the choice stochasticity magnitude effect declines with reward variance (under the same conditions on reward and delay).

Finally, we can examine what happens to the two magnitude effects when the sensitivity parameter *β* changes:

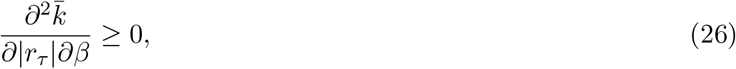

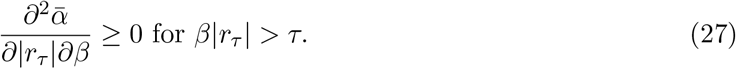

Because 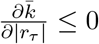, this means that increasing *β* will *decrease* the discounting magnitude effect (i.e., push it closer to 0). This is somewhat counterintuitive, since one might reason that greater sensitivity to reward should translate into a stronger magnitude effect. This intuition is correct for the choice stochasticity magnitude effect: increasing *β* will magnify the dependence of choice stochasticity on reward magnitude. The key implication of this analysis is that a change in sensitivity will push the two magnitude effects in opposite directions.

### Ballard data set description

Ballard and colleagues [18] recruited 1500 subjects for their Study 3. After exclusions, the final sample size was 1382. Subjects considered a hypothetical choice between an immediate reward vs. a reward in one month. Each subject was randomly assigned to one delayed reward magnitude ($20, $50, $100, $200, $2000) and reported the immediate reward that would make them indifferent between the two options. Subjects in the “justification” condition were asked to justify their responses in 2-3 written sentences; subjects in the “no justification” condition did not have to provide any written justification.

### Chavez data set description

Chavez and colleagues [22] collected data from 1284 Mexican students (a mix of high school juniors and seniors and first-year university students). Subjects completed an intertemporal choice questionnaire developed by [44], consisting of 27 questions, each presenting a hypothetical choice between a smaller sooner (immediately available) monetary reward and a later larger one. Monetary amounts were the same as in the original questionnaire but expressed as Mexican pesos rather than U.S. dollars.

### Experimental methods

Two hundred and twenty-one people were recruited from Amazon Mechanical Turk via TurkPrime [45], and paid $1.25 for their participation. To elicit time preferences, we used a choice titration task in which subjects made a series of binary choices between a smaller-sooner reward and a larger-later reward (see, for example, [34]). They faced 40 “titrator” trials, each consisting of 6 hypothetical binary choices between fixed smaller-sooner and larger-later options, distinguished by larger-later delays which varied from 1 to 6 months. The smaller-sooner reward was always $1 in every trial. The larger-later rewards were drawn from a Gaussian distribution with mean $5 truncated to be above $1 and below $9, rounded to the nearest cent. Subjects were randomly assigned to one of two conditions: in the low variance condition, the larger-later distribution had (untruncated) standard deviation 1, while in the high variance condition, the larger-later distribution had (untruncated) standard deviation 5. Empirically, the former condition had variance 1.03 and the latter had variance 4.97. The task was coded in JavaScript using jsPsych [46].

### Model-fitting and comparison

We fit and compared several models with varying degrees of flexibility.

- **QH**: quasi-hyperbolic discounting function, defined by *D_t_>_0_* = *βδ^t^r* and *D_0_* = *r*. We modeled choices using a generalized version of `Eq. 28:

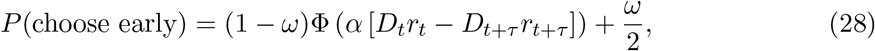

where *α* is the inverse temperature and *ω* is a lapse probability, capturing occasional random responses (see also [24]). All subsequent models share the same choice probability function. This model has 4 free parameters: *β, δ, α, ω*.
- **H0**: standard hyperbolic discounting function, *D_t_* = 1/(1 + *kt*). This model has 3 free parameters: *k, α, ω*.
- **H1**: hyperbolic discounting with baseline- and magnitude-dependent discount factor, using the parametrization of Vincent [24]:

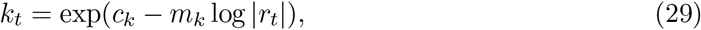

where *c_k_* is a free parameter capturing baseline discounting (i.e., the component of discounting that is independent of magnitude), and *m_k_* captures magnitude-dependent discounting. This model has 4 free parameters: *c_k_*, *m_k_*, *α, ω*.
- **H2**: same as H1, but with baseline- and magnitude-dependent inverse temperature:

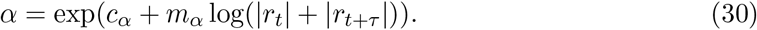 This model has 5 free parameters: *c_k_,m_k_,c_α_,m_α_,ω*.
- **H3**: same as H2, but without the baseline discounting and inverse temperature parameters. In this case, the parametrization simplifies to *k* = |*r_t_*|*^mk^* and *α* = (|*r*_1_| + |*r*_2_|)*^m^*”. This model has 3 free parameters: *m_k_,m_α_,ω*.
- **R1**: hyperbolic discounting with endogenized discount factor, using `Eq. 6, fitting *β* as a free parameter. We approximated 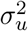 as the empirical variance of the rewards each individual subject observed in the experiment. As in the other models, we use `Eq. 28 to model choices, treating *α* and *ω* as free parameters. This model has 3 free parameters: *β, α, ω*.
- **R2**: hyperbolic discounting with endogenized discount factor and inverse temperature. This model uses the same formulation as R1, but sets *α* using `Eq. 21. The model has 2 free parameters: *β, ω*.

Note that none of the models defined above, except for R1 and R2, are constrained to make the same qualitative predictions as the rational inattention theory. For example, the magnitude scaling parameters in H1-H3 might be 0 on average, or might go in a direction opposite what the theory predicts. Thus, fitting these models gives us the opportunity to test whether the parameter estimates are in qualitative alignment with the rational inattention theory, without making a commitment to the specific parametrization of that theory.

All models were fit using maximum likelihood estimation. To compare models, we computed the *protected exceedance probability* (PXP; [47]), the probability that each model has higher model evidence than all the other models, taking into account the probability that the data may have arisen from a null (chance) model. To approximate model evidence, we used the Bayesian information criterion.

## Data Availability

The data and code for all statistical analyses are available online at https://github.com/sjgershm/rational-discounting.

## Acknowledgments

We are grateful to Xavier Gabaix and David Laibson for helpful discussions. This research was supported by the Office of Naval Research (N00014-17-1-2984), the Center for Brains, Minds and Machines (funded by NSF STC award CCF-1231216), and a research fellowship from the Alfred P. Sloan Foundation.

